# Sterility-Independent Enhancement of Proteasome Function via Floxuridine-Triggered Detoxification in *C. elegans*

**DOI:** 10.1101/2023.11.11.566706

**Authors:** Abhishek Anil Dubey, Natalia A. Szulc, Małgorzata Piechota, Remigiusz A. Serwa, Wojciech Pokrzywa

## Abstract

The ubiquitin-proteasome system (UPS) functionality is vital for proteostasis, contributing to stress resilience, lifespan, and thermal adaptability. In *Caenorhabditis elegans*, proteasome constituents such as the RPN-6 and PBS-6 subunits or the PSME-3 activator are respectively linked to heat resistance, survival at low temperatures (4°C), and longevity at moderate cold (15°C). Since the inhibition of germline stem cells proliferation is associated with robust proteostasis in worms, we utilized floxuridine (FUdR), a compound known for inducing sterility, to examine whether it could reinforce UPS under proteasome dysfunction, particularly to foster cold survival. We demonstrate that FUdR promotes proteasome resilience during its inhibition or subunit deficits, supporting normal lifespan and facilitating adaptation to cold. FUdR’s elevation of the UPS activity occurs independently of main proteostasis regulators and is partly driven by SKN-1-regulated transcription, especially under reduced proteasome function. Additionally, we uncover a FUdR-stimulated detoxification pathway, distinct from both SKN-1 and the germline, with GST-24 emerging as a critical mediator of the UPS buffering. This research underscores FUdR’s role in the UPS modulation and its contribution to survival of worms in low-temperature conditions.

**HIGHLIGHTS:**

- Floxuridine (FUdR) enhances ubiquitin-proteasome system activity in *C. elegans*, independent of primary proteostasis regulators.
- FUdR permits worms to maintain a normal lifespan and facilitates their adaptation to cold in the context of proteasome deficits.
- Acting independently of the germline and SKN-1, FUdR triggers a detoxification pathway, with GST-24 as a pivotal component in modulating the ubiquitin-proteasome system.

## INTRODUCTION

Maintaining protein homeostasis (proteostasis) is pivotal for cellular health, influencing longevity, metabolism, and stress resistance, as exemplified in notable studies on the nematode *Caenorhabditis elegans* (1, 2, 3). A compromised proteostasis system can lead to premature aging, elevated stress sensitivity, and protein-misfolding diseases (2, 3, 4). A growing body of research highlights that reproductive capacity, influenced by signals from proliferating germline stem cells (GSCs), is a critical regulator of proteostasis, impacting metabolic processes in the soma (5). In nematodes, the GLP-1/Notch signaling pathway is crucial for regulating the GSCs pool and establishing germline polarity; moreover, mutations in GLP-1 result in germline loss (6, 7). Remarkably, *glp-1* mutant nematodes exhibit augmented stress resilience, autophagic processes, ubiquitin-proteasome system (UPS) activity, and an extended lifespan. These phenotypic changes are modulated by pivotal molecular regulators, including SKN-1, DAF-12, DAF-16, HSF-1, and TOR (8, 9, 10, 11, 12, 13, 14, 15).

Sterility in *C. elegans* can be also chemically induced using floxuridine (FUdR), a well-documented thymidylate synthase inhibitor and anti-cancer agent. FUdR disrupts DNA and RNA synthesis, causing mitotic cell death and inhibition of protein production (16). Given that adult nematode cells are largely post-mitotic, administering FUdR just prior to sexual maturity primarily prevents progeny development without posing a major effect on the adults. While FUdR does not prolong the lifespan of wild-type worms (16), its introduction at the L4 larval stage extends the lifespans of fat-storing *tub-1* and *gas-1* mutants, which exhibit compromised mitochondrial complex functions (17, 18). Interestingly, FUdR-induced sterility in wild-type adult worms significantly enhances their proteostasis, protein folding, and stress responses (19, 20, 21, 22). While commonly attributed to its sterilizing capabilities, FUdR has been shown to enhance longevity and proteostasis in nematodes devoid of a germline, suggesting that its beneficial effects are not exclusively linked to reproductive inhibition (22). It is postulated that the mechanism for FUdR proteostasis improvements stems from metabolic shifts or the activation of the DNA damage response pathway (23, 24, 25). Nevertheless, the exact mechanism underlying the proteostasis improvements remains elusive.

While proteostasis has traditionally been studied in the context of heat stress, its role under cold has been less explored. Emerging evidence indicates that moderately low temperature (15°C) promote *C. elegans* longevity, enhance proteostasis, and mitigate the accumulation of pathogenic proteins. This protective mechanism is driven predominantly by the PA28γ/PSME-3-activated proteasomes (26). In parallel, Habacher and colleagues, through an RNAi screen, pinpointed PBS-6, a proteasome subunit, as instrumental in cold (4°C) survival (27). This underscores the integral role of the proteasome within the broader proteostasis context during cold adaptation. Building on this foundation, our research sought to discern the relationship between FUdR treatment and its potential role in reinforcing proteostasis, especially when confronting proteasome dysfunction in cold conditions.

Our findings highlight that while proteasome dysfunctions impede the nematode’s cold (4°C) adaptation, FudR treatment enhances their cold tolerance, bolstering proteasome function even in the absence of its subunits. FudR increases proteasome activity and promotes UPS efficiency independently of well-known proteostasis enhancers like RPN-6.1, HSF-1, and DAF-16 (20, 21, 28), but partially relies on SKN-1 regulated transcription. We also reveal that FUdR activates a detoxification pathway distinct from SKN-1 and the germline, with the GST-24 protein emerging as a crucial factor in this enhanced UPS activity. In conclusion, our investigation provides an advanced understanding of the pathways, both sterility-associated and independent, that are modulated by FUdR.

## RESULTS

### Enhancement of proteasome activity via FUdR treatment

To study the impact of FUdR on the *in vivo* UPS activity in *C. elegans,* we utilized a non-cleavable ubiquitin (Ub G76V) fused to a green fluorescent protein (UbV::GFP) under the *sur-5* promoter that remains active in most tissues, as a model to assess the functioning of degradation pathway (29). In this system, the ubiquitin moiety of the GFP substrate is poly-ubiquitylated, leading to its recognition and subsequent degradation by the 26S proteasome. In turn, disruption in the UPS activity leads to an increase in UbV-GFP levels, which can be visualized by fluorescence microscopy or western blotting (30, 31) (Fig 1A). Of note, to account for potential variation arising from the transgene expression which might influence the amount of UbV-GFP, worms were also equipped with a mCherry reporter driven by the same *sur-5* promoter (30), and the GFP fluorescent signal was normalized to the mCherry fluorescence.

**Figure 1.**
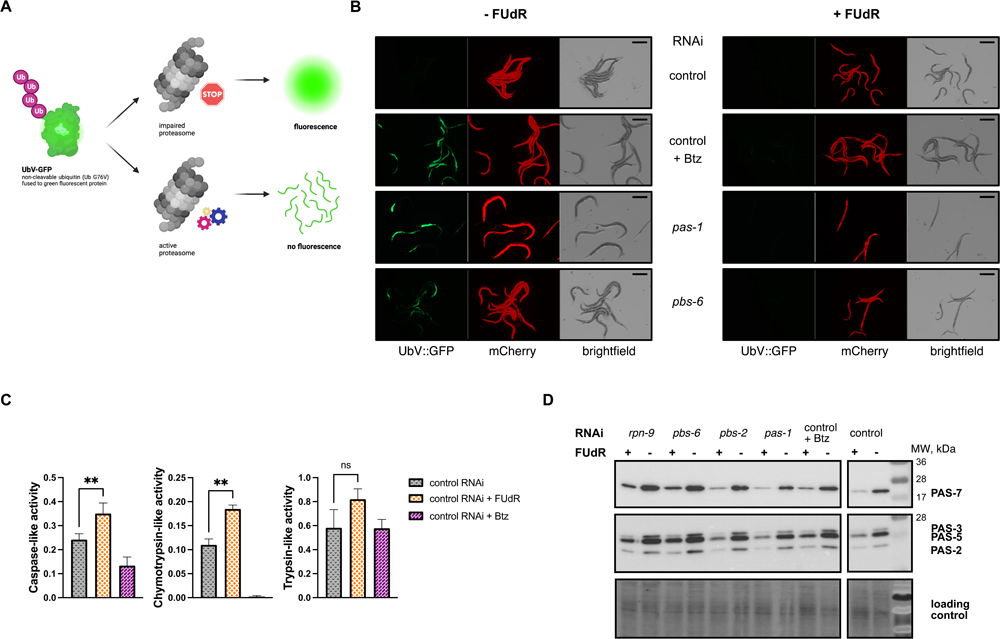
Enhancement of proteasome activity by FUdR. **(A)** A scheme illustrating the method for monitoring *in vivo* UPS activity, employing UbV-GFP as the indicative reporter. **(B)** *In vivo* UPS activity assay showing the effect of FUdR on UbV-GFP turnover in the presence of the proteasome inhibitor bortezomib (Btz) and during the RNAi depletion of *pas-1* and *pbs-6* proteasome subunits. Scale bar corresponds to 400 μm. **(C)** FUdR’s effect on proteasome activity, as measured by trypsin-like, chymotrypsin-like, and caspase-like activity in wild-type worms with or without FUdR treatment. Bortezomib (Btz) served as the negative control. Proteasome activity is represented as slopes obtained from kinetic measurements. The experiments were conducted thrice as separate biological replicates, and significance levels (** - *P* ≤ 0.01) were determined using an unpaired t-test with Welch’s correction. **(D)** Western blot showing levels of PAS-2, PAS-3, PAS-5, and PAS-7 proteasome subunits levels in wild-type worms following the RNAi depletion of RPN-9, PBS-6, PBS-2 and PAS-1 proteasome components in the presence or absence of FUdR, as depicted by using anti-proteasome 20S alpha 1+2+3+5+6+7 antibody. The No-Stain Protein Labeling Reagent was used to confirm equal protein loading.

Our reporter strain was exposed to distinct experimental conditions to elucidate the impact of FUdR on the UPS activity. These conditions were designed to impair proteasome function, either via the direct proteasomal inhibition with bortezomib or through RNA interference (RNAi) targeting of proteasome subunits *pas-1* and *pbs-6*, leading to the accumulation of UbV-GFP. Intriguingly, when FUdR treatment was initiated at the young adult stage, we observed a pronounced increase in the UPS activity across all experimental conditions. This proteostasis rescue effect was evidenced by a concomitant reduction in UbV-GFP accumulation, which is attributable to accelerated GFP degradation. Conversely, worms without FUdR treatment manifested a significant accumulation of UbV-GFP owing to compromised proteasome function (Fig 1B).

Given that FUdR treatment results in sterility in worms, we examined whether the observed effect was a consequence of FUdR-induced sterility. To this end, worms with RNAi-silenced *pbs-6* were subjected to varying concentrations of FUdR and other pyrimidine analogs (32). Our findings revealed that the beneficial impact of FUdR was confined to concentrations that induced nematode sterility (min. 2 μM; Fig S1A), and a similar enhancement in the UPS performance was also observed with other pyrimidine analogs (Fig S1B).

Next, we undertook proteasome activity assays to determine if the observed reduction in UbV-GFP accumulation resulted from elevated proteasome activity. These employed fluorescently labeled probes (28) to measure caspase-like activity (CLA), chymotrypsin-like activity (CTLA), and trypsin-like activity (TLA) in lysates prepared from wild-type worms with or without FUdR treatment. Our data revealed that FUdR was associated with a marked increase in CLA and CTLA activities (Fig 1C). The increase in an activity does not seem to depend on the direct interaction of FUdR with the proteasome, as no changes in the processivity of the recombinant human 26S proteasome were noted in its presence (Fig S1C). To explore the evolutionary conservation of FUdR’s impact on proteasome activity, we observed a slight, non-significant increase in activity in HeLa cells with rising FUdR levels, suggesting the full effect of FUdR might necessitate a multicellular mechanism (Fig S1D).

Next, we conducted a western blot analysis to evaluate if the heightened UPS activity was attributable to increased proteasome subunit levels. This involved analyzing the levels of PAS-{2, 3, 5, 7} in worms treated with FUdR under proteotoxic stress conditions induced either by the presence of bortezomib or upon silencing *pas-1*, *pbs-2*, *pbs-6*, and *rpn-9* proteasome components. Remarkably, we noted that FUdR treatment resulted in the downregulation of PAS-{2, 3, 5, 7} (Fig 1D). This finding mirrors the observation in germline-less worms, known for enhanced proteostasis and downregulation of 26S subunits (28). For this reason, we checked the translation levels in the FUdR-treated nematodes by employing the surface sensing of translation (SUnSET) method which measures puromycin integration into new proteins. We identified a marked decrease in active translation upon FUdR treatment (Fig S1E), which is probably responsible for the observed decrease in proteasome subunit levels (Fig 1D). This effect is likely tied to germline disruption, as similar reduction was evident in germline-less, temperature-sensitive *glp-1(e2144)* mutant worms (Fig S1E). Thus, despite the reduced synthesis of new proteasome subunits, FUdR-treated worms effectively elevated their proteasome activity.

### FUdR bypasses conventional proteostasis regulators

Former studies have underscored the essentiality of a unique proteasome lid subunit, RPN-6.1, in facilitating an upsurge in the UPS activity in sterile *glp-1* mutants (28). Concurrently, sterile worms also necessitate the overexpression of a proteasome core α subunit, PBS-5, to foster proteostasis (9). Because FUdR treatment catalyzes sterility in worms, we conjectured that RPN-6.1 and PBS-5 may be instrumental in the augmentation of proteostasis within FUdR-treated worms.

To validate this hypothesis, we conducted similar as described before *in vivo* assays in the reporter strain, examining UbV-GFP accumulation under conditions involving *glp-1* RNAi treatment, either alone or in conjunction with *pbs-5* or *rpn-6.1* RNAi, and with the variable inclusion of FUdR (Fig 2A). Despite *glp-1* RNAi’s ability to rescue the UbV-GFP accumulation phenotype under PBS-5 depletion conditions, it could not facilitate UbV-GFP turnover without RPN-6.1. In contrast, FUdR treatment succeeded in augmenting UPS activity by remedying UbV-GFP accumulation across all conditions. Further investigations demonstrated that FUdR’s action is not contingent upon any specific proteasome subunit (Fig S2A) nor utilizes autophagy (Fig S2B) to fortify proteostasis. Despite prior assertions that FUdR-mediated resistance to proteotoxic stress is reliant on spermatogenesis (20), our findings suggest that the elevation of UPS activity facilitated by FUdR treatment is not dependent on the spermatogenesis-associated factor, FEM-1 (Fig S2C). This suggests a divergence in the underlying mechanisms responsible for improved proteostasis between *glp-1* dysfunction-induced sterile worms and those rendered sterile by FUdR treatment.

**Figure 2.**
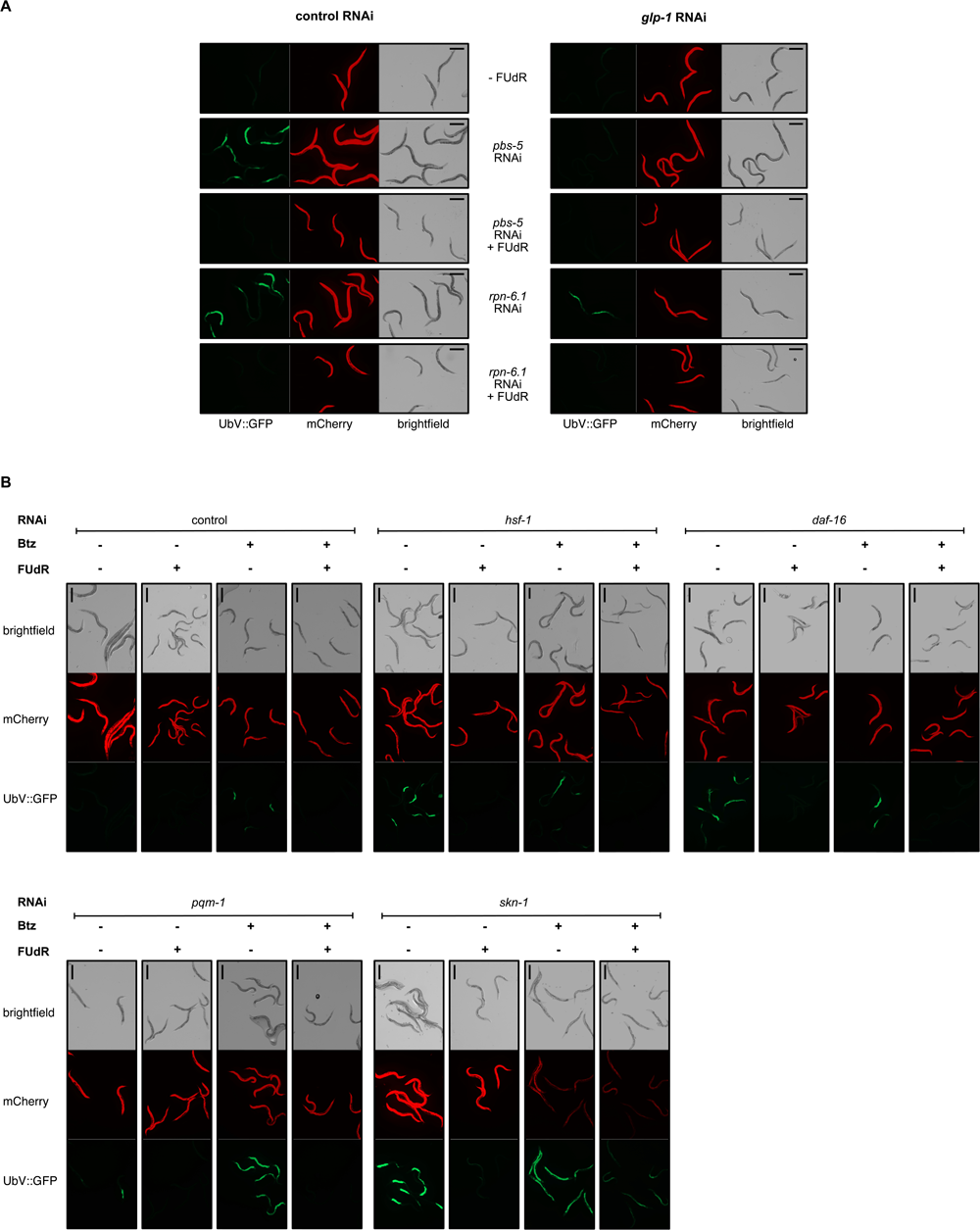
The improved UPS activity mediated by FUdR partially relies on *skn-1*. **(A)** *In vivo* UPS activity assay showing the effect of FUdR on UbV-GFP turnover in worms subjected to *glp-1*, *pbs-5* and *rpn-6.1* RNAi silencing, with or without FUdR. **(B)** *In vivo* UPS activity assay showing the effect of FUdR on UbV-GFP turnover in worms subjected to *hsf-1*, *daf-16*, *pqm-1*, and *skn-1* RNAi silencing in the absence or presence of bortezomib (Btz). In panels A and B, the scale bar corresponds to 400 μm.

To discern the factors essential for UPS activity improvement via FUdR treatment, we induced silencing of *daf-16*, *hsf-1*, *pqm-1*, and *skn-1* factors recognized to regulate proteostasis and confer protection against proteotoxic stress (8, 14, 15, 21, 28), in the presence of bortezomib. FUdR treatment successfully corrected the UbV-GFP accumulation phenotype in cells devoid of HSF-1, DAF-16, and PQM-1. Conversely, the rescue of UbV-GFP accumulation was not completely achieved under *skn-1* RNAi conditions (Fig 2B). This observation aligns with previous research that revealed FUdR-mediated protection against proteotoxic stress independent of *hsf-1* and *daf-16* transcription factors (20).

To corroborate the significance of *skn-1* in the enhancement of UPS activity, we executed a similar as previously described proteasome activity assay, using lysates prepared from wild-type worms under *skn-1* RNAi conditions, with or without FUdR. Our data unveiled that in the absence of SKN-1, worms did not exhibit elevated proteasome activity, even when FUdR was present (Fig S2D). Together, our results outline a mechanistic divergence in enhancing proteostasis in FUdR-treated and sterile *glp-1 C. elegans*, revealing that the effects of FUdR partially rely on SKN-1.

### FUdR prolongs lifespan and promotes cold survival under proteasome deficiencies

The successful induction of hibernation in *C. elegans* under cold conditions is contingent on a β subunit of the proteasome, PBS-6 (27). Also, prior studies have underscored the indispensability of proteasome components for the longevity of worms (33). In the light of our results that highlight FUdR’s capacity to amplify proteasome activity despite its deficits, we conducted cold-survival and lifespan assays (Fig 3A) on wild-type *C. elegans,* with simultaneous RNAi depletion of PAS-1, PBS-2, or PBS-6 proteasome subunits. Our data delineated that FUdR treatment notably increased the lifespan of worms subjected to proteasome disruption (Fig 3B, Fig S3A). Additionally, we discovered that FUdR treatment could effectively re-establish the survival of worms in cold survival assay, even amid proteotoxic stress induced by the depletion of PAS-1, PBS-2, PBS-6, and RPN-9 (Fig 3C). Notably, during cold survival assay, *C. elegans* requires a brief adaptation period at 10°C before exposure to cold incubation at 4°C. The absence of this adaptation period, followed by an abrupt cold incubation for 24 hr or more, has been proven lethal for the worms (34). Hence, we conducted the cold survival assay with or without the adaptation period in the presence of FUdR, under proteasome malfunction caused by knockdown of PBS-6. The FUdR treatment significantly increased worm survival upon sudden cold shock (without an adaptation period), even under compromised proteasome function (Fig S3B).

**Figure 3.**
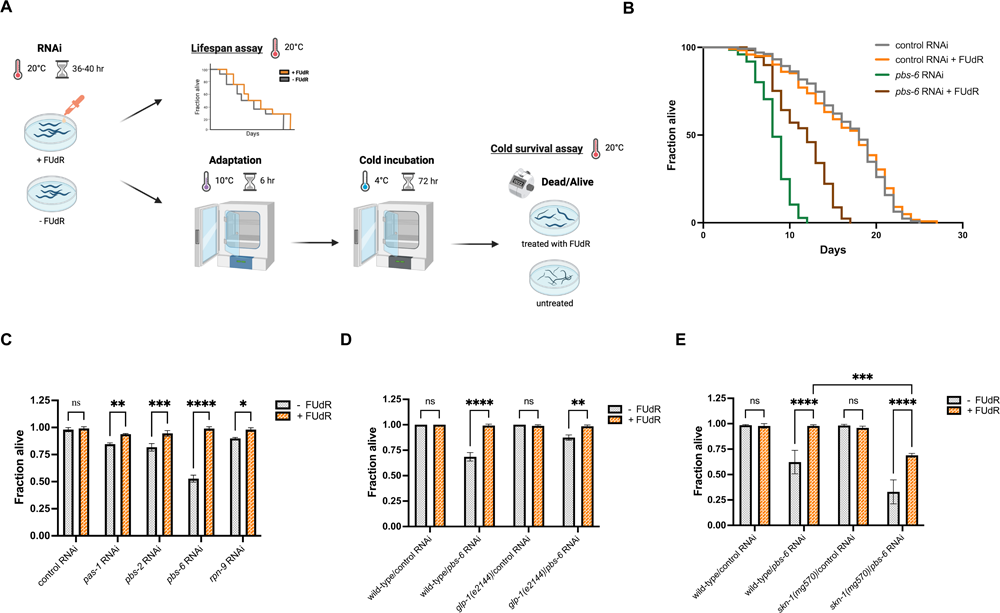
FUdR enhances lifespan and cold survival upon proteasome deficits. **(A)** A schematic representation of cold survival and lifespan assays. **(B)** The lifespan of wild-type worms when exposed to control or *pbs-6* RNAi in the presence or absence FUdR. Number of worms used in the study: control RNAi: n=123 (+ FUdR) or n=139 (-FUdR); *pbs-6* RNAi: n=129 (+ FUdR) or n=149 (-FUdR). **(C)** The impact of FUdR on the cold survival rate of wild-type worms during the knockdown of PAS-1, PBS-2, PBS-6, and RPN-9, considering both FUdR-treated and untreated conditions. Data was analyzed using two-way ANOVA and the significance levels obtained from the Šidák’s multiple comparisons test are indicated for the compared conditions (ns - not significant, * - *P* ≤ 0.05, ** - *P* ≤ 0.01, *** - *P* ≤ 0.001, **** - *P* ≤ 0.0001). **(D)** The impact of FUdR on the cold survival of wild-type and *glp-1(e2144)* worms subjected to control and *pbs-6* RNAi. Data was analyzed using two-way ANOVA and the significance levels obtained from the Šidák’s multiple comparisons test are indicated for the compared conditions (ns - not significant, ** - *P* ≤ 0.01, **** - *P* ≤ 0.0001). **(E)** The impact of FUdR on the cold survival of wild-type and *skn-1(mg570)* worms subjected to control and *pbs-6* RNAi. Data was analyzed using two-way ANOVA and the significance levels obtained from the Tukey’s multiple comparisons test are indicated for the compared conditions (ns - not significant, *** - *P* ≤ 0.001, **** - *P* ≤ 0.0001). In panels C-E, at least 90 animals were scored in three independent biological replicates.

To scrutinize the impact of tissue-specific proteostasis collapse on cold survival, we silenced *pbs-6* in wild-type strains and strains permitting tissue-specific RNAi depletion in muscles, intestines, germline, or neurons, again with or without FUdR. Our findings revealed that the knockdown of PBS-6 in neurons and intestines had the most pronounced impact on cold survival, which can be fully rescued by the FUdR treatment (Fig S3C).

Subsequently, we posited that FUdR-induced sterility plays a role in enhancing cold survival, a hypothesis supported by experiments involving both *glp-1(e2144)* sterile mutants and wild-type worms. The cold survival assay indicated enhanced survival in *glp-1(e2144)* mutants compared to wild-type worms when PBS-6 was depleted (Fig 3D). Moreover, this survival advantage of *glp-1(e2144)* worms was further potentiated in the presence of FUdR, suggesting that the mechanisms of cold tolerance induced by FUdR are likely distinct from the protective effects conferred by sterility alone.

As previously demonstrated, the SKN-1 transcription factor is partially required for FUdR to fully restore the enhanced proteostasis phenotype (Fig 2B). Therefore, we executed a cold survival assay involving both wild-type and *skn-1(mg570)* loss of function mutants, with or without RNAi silencing of *pbs-6* and FUdR. While the *skn-1(mg570)* worms exhibited no notable decline in survival after cold incubation in the control RNAi condition, a significant reduction in survival was noted for *skn-1(mg570)* mutants when subjected to proteasome dysfunction by *pbs-6* silencing and cold exposure (Fig 3E). The observed decline was, albeit not entirely, rescued by FUdR treatment.

### FUdR induces a detoxification pathway buffering UPS during proteasome inhibition

Based on our findings from the SUnSET assay (Fig. S1E), we speculated that the suppressive action of FUdR on translation could play a significant role in supporting UPS activity, given the well-established challenge that newly produced proteins pose to the proteostasis machinery (35, 36, 37, 38). However, despite the pronounced FUdR-mediated reduction in new protein synthesis in *skn-1(mg577)* mutant worms (Fig 4A), the role of the SKN-1 transcription factor remains significant for FUdR’s ability to promote UPS function when confronted with proteasome inhibitors like bortezomib (Fig 2B). Furthermore, our findings show that while both FUdR and the *glp-1(e2144)* mutation significantly reduce translation, FUdR distinctively triggers UPS-enhancing mechanisms in germline-less *glp-1* mutants, evident in its ability to circumvent the requirement for RPN-6.1 (Fig 2A). Prior research has demonstrated that in response to global translation inhibition, ribosome availability facilitates selective translation, especially of stress response factors (37, 38, 39, 40). Given these insights, we hypothesized that the mechanism underlying FUdR’s action could be centered on the selective translation of the UPS buffering proteins, which should also occur in *glp-1* and *skn-1* mutants. To investigate this, we conducted a targeted proteomic analysis of adult wild-type, *glp-1(e2144)*, and *skn-1(mg570)* worms treated with 400 µM FUdR, aiming to identify proteins whose level is uniquely increased by FUdR. We identified and consistently quantified 5855 protein groups across all experimental conditions and three independent biological replicates. The analysis revealed that the FUdR treatment affected total levels of numerous proteins (717 up-and 821 down-regulated in wild-type, 416 up-and 442 down-regulated in *glp-1(e2144)*, 1132 up-and 1061 down-regulated in *skn-1(mg570)* worms) (Table S1).

**Figure 4.**
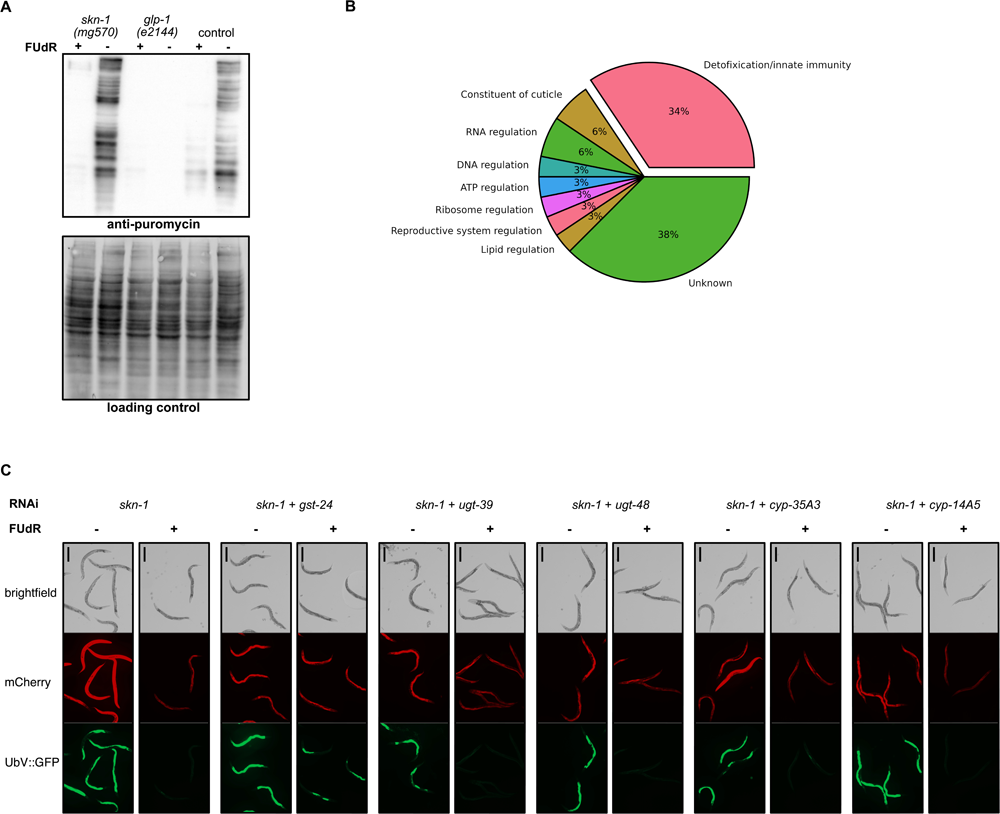
FUdR promotes detoxification pathway independent of *skn-1* and *glp-1.* **(A)** Western blot showing global translation activity in wild-type, *glp-1(e2144)*, and *skn-1(mg570)* worms, in the presence or absence of FUdR, as depicted by using anti-puromycin antibody. The No-Stain Protein Labeling Reagent was used to confirm equal protein loading. **(B)** Pie chart representing the functional categories of proteins from our proteomics study that exhibited a marked increase (fold change > 1.0), observed consistently in wild-type, *glp-1(e2144)*, and *skn-1(mg570)* worms specifically due to FUdR treatment. The significance of proteomic findings was confirmed through student’s t-tests, setting a *P*-value threshold < 0.05 while maintaining the false discovery rate under 0.01. Functional annotations were compiled by manual review. **(C)** *In vivo* UPS activity assay showing the effect of FUdR on UbV-GFP turnover in worms subjected to *skn-1* RNAi silencing alone or in combination with *gst-24*, *ugt-39*, *ugt-48*, *cyp35A3*, and *cyp14A5* RNAi silencing, with or without FUdR. Scale bar corresponds to 400 μm.

To uncover pathways activated solely by FUdR, we manually categorized functions of proteins up-regulated across all three strains upon FUdR treatment (Table S2, Fig S4A). Our analysis revealed that over 33% of these proteins are implicated in detoxification and innate immunity pathways (Fig 4B), indicating that FUdR induces a detoxification response that is autonomous of both germline and *skn-1*. To eliminate the possibility that this induction is due to metabolites produced by bacterial metabolism of FUdR or bacterial infection, we examined UbV-GFP reporter degradation in worms fed with both live and killed HT115 *Escherichia coli*, in the presence or absence of bortezomib and FUdR. Our results demonstrated that FUdR effectively mitigated UbV-GFP accumulation under both bacterial conditions (Fig S4B).

To further elucidate our proteomic findings and clarify if detoxification proteins are involved in FUdR-mediated proteostasis enhancement, we focused on RNAi-mediated depletion of the most abundant detoxifying proteins - specifically GST-24, UGT-39, UGT-48, CYP-35A3, and CYP-14A5 - that were uniquely upregulated under FUdR treatment, while monitoring the turnover of the UBV-GFP reporter. Our data showed that GST-24 and CYP-14A5, when depleted individually, can affect UPS activity (Fig S4C). Notably, FUdR effectively reinstates proteasomal degradation in worms where SKN-1 is silenced alone or combined with UGT-39, UGT-48, CYP-35A3, or CYP14A5 (Fig 4C). This restorative effect is not observed when GST-24 is co-depleted with SKN-1 (Fig 4C), highlighting crucial role of GST-24 in the detoxification pathway that contributes to FUdR’s enhancement of the UPS. However, the singular RNAi depletion of GST-24 from the array of FUdR-upregulated detoxifying proteins did not compromise the safeguarding effect of FUdR on the worms’ resistance to low temperatures upon PBS-6 depletion (Fig S4D), indicating the cumulative action of other mechanisms.

A previous study has suggested that dopamine signaling induces the xenobiotic stress response and improves proteostasis (41). To validate whether the neuronal signal is essential in improving the UPS activity upon FUdR induction, we measured the turnover of UbV-GFP reporter upon depleting the sensory neuron ciliary components CHE-12 and CHE-13 (42, 43) with or without knockdown of SKN-1. Our findings indicate that individual silencing of either *che-12* or *che-13* exerts no discernible impact on UPS activity (Fig S4E). Upon simultaneous silencing of SKN1 and either CHE-12 or CHE-13, we noticed an alteration in proteostasis, further emphasized by FUdR’s inability to fully ameliorate UbV-GFP degradation when CHE-12 or CHE-13 was explicitly targeted (Fig S4E). This suggests the involvement of neuronal signaling in the regulation of proteostasis by FUdR.

## DISCUSSION

While FUdR is primarily known for inducing reproductive arrest, it also exerts secondary effects on various facets of *C. elegans* physiology, such as proteostasis and stress response mechanisms (16, 19, 20, 21, 22). Our data indicate that FUdR augments the proteasome’s caspase-like and chymotrypsin-like activities, akin to the effects observed in the sterile *glp-1(e2144)* mutant (28). Although both FUdR and germline-deficient worms are known to elevate proteasome activity and stress resistance (13, 22), FUdR appears to act independently of germline signaling. Enhanced proteostasis in *glp-1* mutants is contingent upon RPN-6.1, a lid subunit of the 26S proteasome (28), whereas the effect of FUdR is not. Moreover, these mechanisms further diverge: while *glp-1* mutants are dependent on *daf-16* for both lifespan extension and elevated proteasome activity (28), the enhancement of UPS activity in FUdR-treated worms is partially attributable to the *skn-1* transcription factor, but not *daf-16*.

Our findings demonstrate that FUdR treatment initiates a xenobiotic detoxification pathway involving innate immune response proteins independently of SKN-1 and germline signaling. Although SKN-1 generally activates proteasome subunits and is influenced by lipids in sterile animals (14, 44), FUdR does not rely on specific 26S proteasome subunits or autophagy for its effects. Instead, SKN-1 appears to have an alternative role in enhancing UPS activity in the presence of FUdR, possibly through SKN-1-dependent and independent xenobiotic pathways (45, 46, 47). We note that the latter is involved in the up-regulation of i.a., GST-24, previously linked to stress resistance in *C. elegans* (48). Furthermore, given that FUdR induces DNA breaks, activating damage repair pathways (24), and that DNA breaks in the germline improve UPS activity through an innate immune response (49), it is plausible that FUdR triggers this detoxification pathway in response to DNA damage. Parallelly, we observed that treatment with FUdR enhances the ability of *C. elegans* to withstand cold stress at 4°C, even bypassing the usual need for a preparatory acclimatization phase at 10°C. This raises the possibility that FUdR treatment mimics or induces adaptive processes normally activated during this acclimatization period. Previous research has shown that moderate cold exposure at 10°C triggers shifts in lipid metabolism, which are likely crucial for enhanced cold resistance (50, 51). Interestingly, lipid metabolism in *C. elegans* is deeply interwoven with innate immune responses; specific fatty acids and their synthesizing enzymes are essential for immune gene expression and pathogen resistance (52). Given that FUdR pre-treatment activates detoxification pathways in the worms prior to exposure to severe cold, it is conceivable that such pathways and lipid metabolism may interact to confer cold resilience.

It is noteworthy that the beneficial effects of FUdR in promoting worms survival and protein turnover are not dependent on evolutionarily preserved transcriptional regulators DAF-16 and PQM-1, recognized for their cooperative role in enhancing resilience to cold conditions through the upregulation of FTN-1/ferritin (53). This delineates a distinct mechanistic route through which FUdR modulates cellular homeostasis and also raises intriguing questions about the potential cross-talk between various stress response pathways and how they might be selectively activated or bypassed.

Our study offers a comprehensive analysis of the diverse functional roles of FUdR in *C. elegans*, emphasizing its modulation of the UPS efficiency, detoxification pathways, and cold resilience mechanisms. Although there are marked differences in the architecture of immune systems between nematodes and higher organisms, several innate immunity and detoxification mechanisms are evolutionarily preserved, suggesting potential translational relevance (54). These findings thus warrant further exploration of the influence of FUdR on proteasomal activity, particularly in the context of its co-administration with agents like bortezomib in oncological patients (55, 56). Additionally, it would be valuable to study whether FUdR or related molecules have the potential to enhance cellular resilience in mammals subjected to environmental stressors, including temperature fluctuations. Such research could extend our grasp of the adaptability-promoting properties of these compounds across different species.

## MATERIALS AND METHODS

### Worm maintenance

*C. elegans* were cultured on nematode growth medium (NGM) plates seeded with OP50 or HT115 *E. coli* strains. These culture conditions were maintained at various temperatures, namely 16°C, 20°C, or 25°C, depending on both the worm strain and the specific requirements of the experimental protocol, following standard *C. elegans* culture techniques (57). Dead bacterial food sources were prepared by exposing bacterial cultures to paraformaldehyde (58). Worm strains utilized for different experimental setups are cataloged in Table 1. Unless stated otherwise, NGM plates were supplemented with 400 μM of floxuridine (FUdR; Cat: F0503; Sigma Aldrich). In some instances, other pyrimidine analogs were employed, including 5-fluorocytosine (FC; Cat: 543020; Sigma Aldrich), 5-fluorouracil (FU; Cat: F6627; Sigma Aldrich), and 5-fluoro-2′-deoxycytidine (FCdR; Cat: F5307; Sigma Aldrich), as well as proteasome inhibitor bortezomib (Cat: 504314; Sigma Aldrich), each at a concentration of 10 μM.

**Table 1.** *C. elegans* strains used in this study.

| Strain | Source | Purpose |
| --- | --- | --- |
| *C. elegans*: Bristol (N2) | CGC<sup>1</sup> | Wild-type; control worms |
| *skn-1(mg570)* | CGC | *skn-1* mutant |
| *glp-1(e2144)* | CGC | *glp-1* mutant |
| hhls72[*unc-119(+); sur-5::mCherry*],<br>hhls64[*unc-119(+); sur-5::UbiV-GFP*] | (30) | UPS activity reporter |
| *rde-1(mkc36); mkcSi13* | (59) | Germline-specific RNAi |
| *rde-1(ne219); kzlS20* | (60) | Body wall muscle-specific RNAi |
| *rde-1(ne219); kbls7* | (61) | Intestine-specific RNAi |
| TU3311 [*uls60 (unc-119p::YFP + unc-119p::sid-1)*] | (62) | Neuron-specific RNAi |

<sup>1</sup> Caenorhabditis Genetics Center

### RNA interference

Gene silencing was carried out by the standard RNAi feeding method using clones from the Ahringer *C. elegans* RNAi library (63). RNAi was applied at different stages for each target gene, as described in Table S3. NGM plates supplemented with 1 mM IPTG (Cat: IPT001; BioShop) and 25 μg/μl carbenicillin (Art. No. 6344.2; Carl Roth Gmbh & Co.) were seeded with bacteria expressing double-stranded RNA from L4440 plasmid; *E. coli* HT115 (DE3) bacteria containing empty L4440 plasmid were used as control.

### Lifespan assay

Synchronized late-stage L4 worms were silenced for *pbs-6*, *pas-1*, *pbs-2,* or control RNAi (empty vector) for 40 hr with or without FUdR. Subsequently, lifespan measurements were carried out at a constant temperature of 20°C. Approximately 30 nematodes were maintained on each 6 cm diameter agar plate for the duration of the lifespan assay. Daily evaluations were performed to monitor the worms for movement and pharyngeal pumping as vitality indicators. Worms were deemed to have reached the end of their lifespan if they failed to display these physiological activities. Any animals manifesting bagging phenotypes or found to have crawled off the agar surface were censored. Data was analyzed using the Log-Rank (Mantel-Cox) test in the GraphPad Prism 9 software. The experiments were not randomized. No statistical methods were used to predetermine the sample size. The investigators were blinded to allocation during experiments.

### Cold survival assay

To assess the impact of specific gene knockdown and FUdR treatment on cold tolerance, approximately 30 late-stage L4 *C. elegans* were subjected to gene silencing. The gene silencing was performed for 40 hr at a controlled temperature of 20°C, in the presence or absence of FUdR, according to protocols detailed in the study by Habacher and colleagues (27).

#### Adaptation and exposure to cold conditions

Following the gene silencing phase, the nematodes were acclimatized to a moderately cold environment at 10°C for 6 hr. Subsequently, the worms were exposed to a colder temperature setting at 4°C and maintained at this condition for 72 hr.

#### Recovery and scoring

After the 3-day cold exposure, the worms were allowed to recover at 20°C, lasting between 3 and 4 hr. Following the recovery phase, an assessment was carried out to distinguish between live and dead specimens.

#### Cold incubation treatment

As a control, cold incubation was administered by directly transferring the worms from a 20°C environment to a 4°C setting for 72 hr without prior acclimatization.

### UbV-GFP substrate turnover assay

To explore the turnover dynamics of the ubiquitin-based substrate in the presence of gene knockdown and various pharmacological treatments, we utilized L4 stage hermaphrodites of two specific strains: hhls72*[unc-119(+); sur-5::mCherry]* and hhls64*[unc-119(+); sur-5::UbiV-GFP]*. Nematodes were silenced for the target genes over 40 hr at a consistent temperature of 20°C. During this gene-silencing stage, the worms were treated with varying concentration of FUdR, FU, FC, FCdR, and 100 nM bortezomib. Post-treatment, nematodes were subjected to fluorescence imaging employing an Axio Zoom.V16 microscope outfitted with an Axiocam 705 monochrome CMOS camera (Carl Zeiss). Images were captured across brightfield, green, and red channels. Subsequent data processing was performed using the AxioVision 4.7 analysis software (Carl Zeiss) to quantify the rates of substrate turnover, as reflected by the intensity and distribution of fluorescence signals.

### Proteasome activity measurement in worms

Approximately 7000 L4 stage worms were silenced for the respective genes at a concentration of 400 μM for 40 hr at 20°C. Next, worms were harvested and lysed in 50 mM Tris-HCl, 250 mM sucrose, 5 mM MgCl_2_, 0.5 mM EDTA, 2 mM ATP, and 1 mM dithiothreitol at pH 7.5. Bortezomib was introduced to the lysate as a control at a concentration of 20 nM. The caspase-like, chymotrypsin-like, and trypsin-like activities of the proteasome were subsequently measured following the methods described (28). Proteasome activity was also measured using 5 nM of recombinant human 26S proteasome (Cat: E-365; R&D Systems).

### Proteasome activity measurement in cells

HeLa Flp-In T-Rex cell line (a kind gift from R. Szczęsny) was cultured in Dulbecco’s Modified Eagle’s Medium (Cat: 6429; Sigma Aldrich) supplemented with 10% fetal bovine serum (Cat: F9965; Sigma Aldrich) and 1% Antibiotic-Antimycotic (Cat: 15240096; Gibco) at 37°C with 5% CO_2_ in a humidified incubator. HeLa cells were seeded in black 96-well tissue culture plates (Cat: 655090; Greiner) at a density of 20,000 cells in a total volume of 100 µl per well. The next day, the cells were subjected to a 6-hr treatment with FUdR at 2.5 µg/ml concentration in the growing media. Where indicated, the cells were also treated with 10 nM bortezomib (Cat: 504314; Merck). The control cells received dimethyl sulfoxide. Using the Proteasome 20S Activity Assay Kit (Cat: MAK172; Sigma Aldrich), 10.75 µl of LLVY-R110 Substrate was mixed with 4300 µl of the Assay Solution to create the Proteasome Assay Loading Solution, according to the manufacturer’s guidelines. Subsequently, 100 µl of this prepared solution was accurately pipetted into each assay plate well. The plate was then incubated at 37°C for 2 hr. After the incubation period, the fluorescence intensity (λ_ex_ = 490 nm, λ_em_ = 525 nm) was gauged using the TECAN Infinite 200 Pro plate reader equipped with the Magellan Pro software to ascertain proteasome activity.

### Surface sensing of translation assay

The surface sensing of translation (SUnSET) assay was employed to evaluate protein synthesis rates, following the delineated methodology (64). A minimum of 7,000 worms were incubated in 4 mL of S-complete liquid media supplemented with 750 μL of 10X *E. coli* food, engineered to express double-stranded RNA against targeted genes, and 0.5 mg/mL puromycin (Cat: ant-pr-1; Invivogen). The incubation occurred at 20°C with a consistent shaking speed of 200 rpm for 5 hr. The worms were then harvested, washed three times in M9 buffer, and subsequently lysed in a lysis buffer (50 mM KCl, 10 mM Tris-HCl pH 8.2, 2.5 mM MgCl_2_, 0.07% NP-40, 0.7% Tween-20, 0.1% gelatin), supplemented with protease inhibitor (Cat: 04693159001; Roche). Protein concentration was determined using the Pierce Rapid Gold BCA Protein Assay Kit (Cat: A53225; Thermo Fisher Scientific) and western blotting.

### Western blotting

Proteins were separated by electrophoresis on 12% acrylamide gels using a running buffer containing 25 mM Tris, 190 mM glycine, and 0.1% SDS. Electrophoresis was conducted at a constant voltage of 150 V. Subsequently, protein samples were transferred to PVDF membranes through a wet transfer method at 100 V for one hour. The transfer buffer contained 25 mM Tris, 190 mM glycine, and 10% methanol, with a pH of 8.3. For protein visualization, the membranes were treated with Invitrogen No-Stain Protein Labeling Reagent (Cat: A44717; Thermo Scientific) according to the manufacturer’s guidelines. Following this, the membranes were blocked in a solution of 5% skimmed milk dissolved in TBST buffer (50 mM Tris, 150 mM NaCl, 0.1% Tween 20, pH 7.5) for 45 min at room temperature. Primary antibody incubation was performed overnight at 4°C using either anti-GFP antibody (Cat: GF208R; Thermo Fisher Scientific), anti-PAS-7 antibody (Cat: CePAS7; Developmental Studies Hybridoma Bank (DSHB)), anti-proteasome 20S alpha 1+2+3+5+6+7 antibody (Cat: ab22674; Abcam) or anti-puromycin (Cat: MABE343; Merck), both prepared in 5% skimmed milk in TBST buffer. Post-incubation, membranes were washed three times with TBST for 10 min each and incubated with the appropriate secondary antibodies, prepared in the blocking solution, for 45 min at room temperature. Membranes were subsequently developed and visualized using a ChemiDoc Imaging System from Bio-Rad.

### Proteomics analysis

*C. elegans* were extracted using the Sample Preparation by Easy Extraction and Digestion (SPEED) protocol (65). In brief, *C. elegans* were solubilized in concentrated Trifluoroacetic Acid (TFA; Cat: T6508; Sigma Aldrich) (cell pellet/TFA 1:2-1:4 (v/v)) and incubated for 2-10 min at room temperature. Next, samples were neutralized with 2 M Tris-Base buffer using 10x volume of TFA and further incubated at 95°C for 5 min after adding Tris(2-carboxyethyl)phosphine (final concentration 10 mM) and 2-chloroacetamide (final concentration 40 mM). Turbidity measurements determined protein concentrations at 360 nm, adjusted to the same concentration using a sample dilution buffer (2M TrisBase/TFA 10:1 (v/v)), and then diluted 1:4-1:5 with water. Digestion was carried out overnight at 37°C using trypsin at a protein/enzyme ratio of 100:1. TFA was added to a final concentration of 2% to stop digestion. The resulting peptides were labeled using an on-column TMT labeling protoco (66). TMT-labeled samples were compiled into a single TMT sample and concentrated.

Peptides in the compiled sample were fractionated (8 fractions) using the Pierce High pH Reversed-Phase Peptide Fractionation Kit (Cat: 84868; Thermo Fisher Scientific). Prior to liquid chromatography–mass spectrometry (LC-MS) measurement, the peptide fractions were reconstituted in 0.1% TFA, 2% acetonitrile in water. Chromatographic separation was performed on an Easy-Spray Acclaim PepMap column 50 cm long × 75 µm inner diameter (Cat: PN ES903; Thermo Fisher Scientific) at 55°C by applying 120 min acetonitrile gradients in 0.1% aqueous formic acid at a flow rate of 300 nl/min. An UltiMate 3000 nano-LC system was coupled to a Q Exactive HF-X mass spectrometer via an easy-spray source (all Thermo Fisher Scientific). The Q Exactive HF-X was operated in TMT mode with survey scans acquired at a resolution of 60,000 at m/z 200. Up to 15 of the most abundant isotope patterns with charges 2-5 from the survey scan were selected with an isolation window of 0.7 m/z and fragmented by higher-energy collision dissociation (HCD) with normalized collision energies of 32, while the dynamic exclusion was set to 35 s. The maximum ion injection times for the survey and tandem mass spectrometry (MS/MS) scans (acquired with a resolution of 45,000 at m/z 200) were 50 and 96 ms, respectively. The ion target value for MS was set to 3e6 and for MS/MS to 1e5, and the minimum AGC target was set to 1e3.

The data were processed with MaxQuant v. 1.6.17.0 (67), and the peptides were identified from the MS/MS spectra searched against Uniprot *C. elegans* reference proteome (UP000001940) using the built-in Andromeda search engine. Raw files from the liquid chromatography with tandem mass spectrometry (LC-MS/MS) measurements of 8 tryptic peptide fractions were analyzed together. Reporter ion MS2-based quantification was applied with reporter mass tolerance = 0.003 Da and min. reporter PIF = 0.75. Cysteine carbamidomethylation was set as a fixed modification, and methionine oxidation, glutamine/asparagine deamination, and protein N-terminal acetylation were set as variable modifications. For *in silico* digests of the reference proteome, cleavages of arginine or lysine followed by any amino acid were allowed (trypsin/P), and up to two missed cleavages were allowed. The false discovery rate (FDR) was set to 0.01 for peptides, proteins, and sites. A match between runs was enabled. Other parameters were used as pre-set in the software. Unique and razor peptides were used for quantification, enabling protein grouping (razor peptides are the peptides uniquely assigned to protein groups and not to individual proteins). Reporter intensity corrected values for protein groups were loaded into Perseus v. 1.6.10.0. (68). Standard filtering steps were applied to clean up the dataset: reverse (matched to decoy database), only identified by site, and potential contaminant (from a list of commonly occurring contaminants included in MaxQuant) protein groups were removed. Reporter intensity corrected values were log2 transformed, and protein groups with all values were kept. Reporter intensity values were then normalized by median subtraction within TMT channels. Student’s t-tests (permutation-based FDR = 0.001, S0 = 0.1) were performed on the dataset to return protein groups, whose levels were statistically significantly changed between the sample groups investigated.

## DATA AVAILABILITY

The proteomics data was uploaded to the ProteomeXchange Consortium (69) via the PRIDE partner repository (70) with the dataset identifier PXD045805. The datasets used and/or analyzed during the current study are available from the corresponding author upon reasonable request.

## AUTHOR CONTRIBUTIONS

**AAD**: Data curation; conceptualization; formal analysis; investigation; visualization; writing. **NAS**: Data curation; formal analysis; visualization; writing. **MP**: Investigation. **RAS**: Investigation (proteomics). **WP**: Conceptualization; resources; supervision; funding acquisition; validation; project administration; writing. All authors read and approved the manuscript.

## ACKNOWLEDGMENTS

Proteomic measurements were performed at the Proteomics Core Facility, IMol Polish Academy of Sciences. Special thanks to Dorota Stadnik for help with sample preparation and LC-MS/MS measurements. We thank the Caenorhabditis Genetics Center (funded by the NIH National Center for Research Resources, P40 OD010440) for *C. elegans* strains. We thank Roman Szczęsny for the HeLa Flp-In T-Rex cell line. We also acknowledge Marta Niklewicz and Karolina Milcz for their technical assistance and Rafał Ciosk for comments on the manuscript.

## SUPPLEMENTARY TABLES’ AND FIGURES’ LEGENDS

**Table S1.** Results of proteomics analysis showing changes in protein abundance in wild-type, *glp-1(e2144)*, and *skn-1(mg570)* worms in the presence or absence of FUdR.

**Table S2.** Proteins up-regulated in wild-type, *glp-1(e2144)*, and *skn-1(mg570)* upon FUdR treatment with manually categorized functions.

**Table S3.** Worm stages designated for specific RNAi treatments.

**Figure S1.**
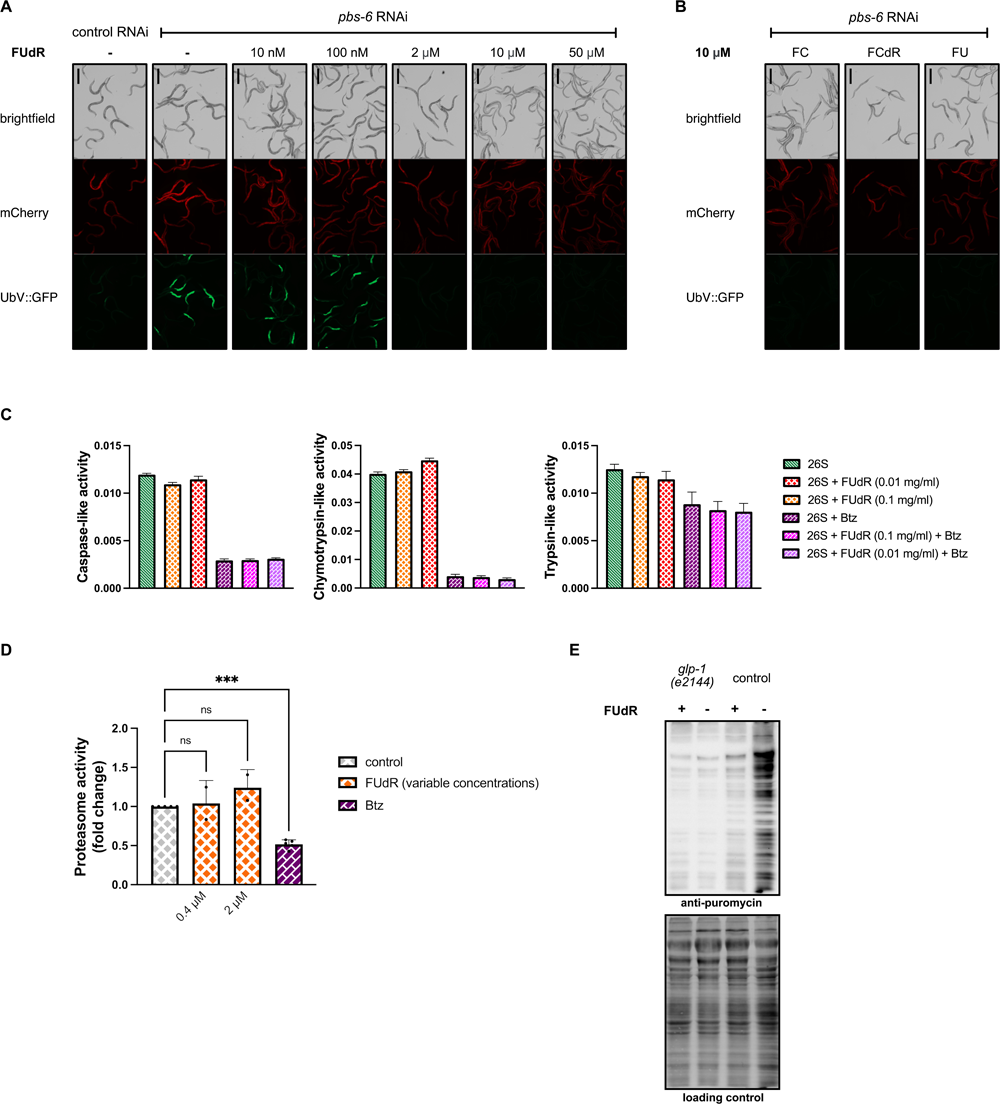
Pyrimidine analogs improve UPS activity. **(A)** *In vivo* UPS activity assay showing the effect of different concentrations of FUdR on UbV-GFP turnover. **(B)** *In vivo* UPS activity assay showing the effect of different pyrimidine analogs, 5-fluorouracil (FU), 5-fluorocytosine (FC), and 5-fluorodeoxycytidine (FCdR), on UbV-GFP turnover. In panels A and B, the scale bar corresponds to 400 μm. **(C)** Effect of different concentrations of FUdR on the activity of purified human 26S proteasome in HeLa cells, as measured by trypsin-like, chymotrypsin-like, and caspase-like activity in wild-type worms with or without FUdR treatment. Bortezomib (Btz) served as the negative control. Proteasome activity is represented as slopes obtained from kinetic measurements. The experiments were conducted thrice as separate biological replicates. **(D)** FUdR effect on chymotrypsin-like proteasome activity in HeLa cells. Cells were treated with final concentrations of 0.4 and 2 µM of FUdR and 10 nM bortezomib (Btz) as control for 6 hr. The assay was conducted by incubating the cells with 100 µl Proteasome Assay Loading Solution for 2 h as described in the methods. Results from three technical replicates were corrected for background by subtracting the fluorescence of the medium without cells and further normalized to dimethyl sulfoxide control. The graph shows the average values obtained from either two or four biological replicates for experiments that involve FUdR or Btz, respectively. **(E)** Western blot showing global translation activity in wild-type and *glp-1(e2144)* worms, in the presence or absence of FUdR, as depicted by using the anti-puromycin antibody. The No-Stain Protein Labeling Reagent was used to confirm equal protein loading.

**Figure S2.**
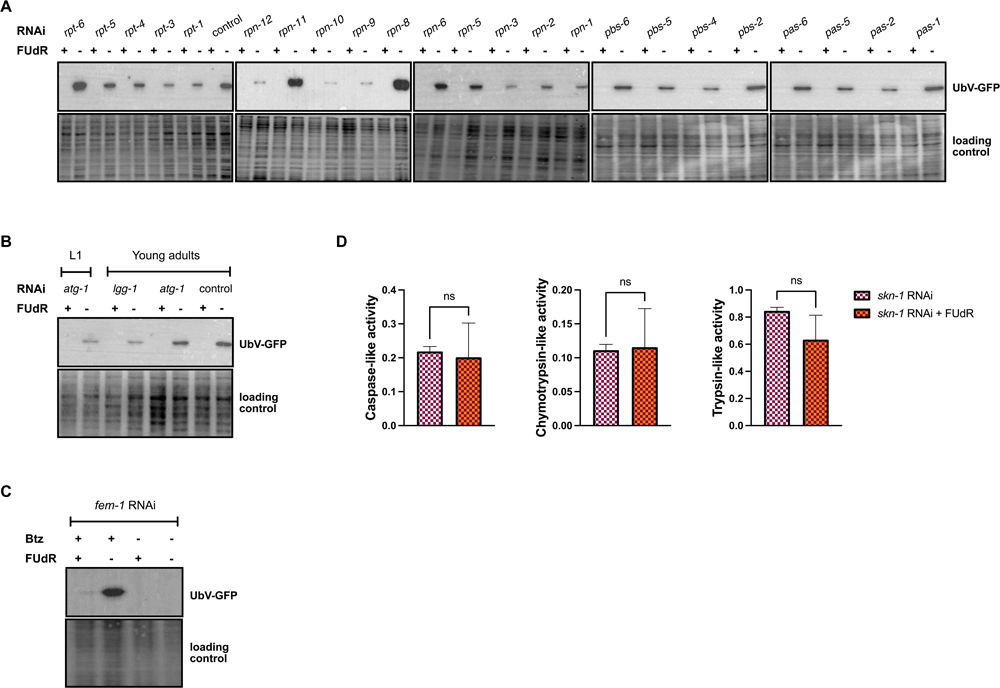
FUdR buffers UPS activity under proteasome-compromised conditions. **(A)** Western blot showing the impact of FUdR on UbV-GFP reporter turnover upon depletion of various 26S subunits in control worms, as depicted by using anti-GFP antibody. The No-Stain Protein Labeling Reagent was used to confirm equal protein loading. **(B)** Western blot showing degradation of UbV-GFP in control, *atg-1* RNAi (applied at either L1 or young adult stages), and *lgg-1* RNAi (applied at the young adult stage) worms co-treated with bortezomib (Btz) in the presence or absence of FUdR, as depicted by using anti-GFP antibody. The No-Stain Protein Labeling Reagent was used to confirm equal protein loading. **(C)** Western blot showing degradation of UbV-GFP upon RNAi depletion of *fem-1* in control worms, with or without bortezomib (Btz) or FUdR, as depicted by using anti-GFP antibody. The No-Stain Protein Labeling Reagent was used to confirm equal protein loading. (**D)** FUdR’s effect on proteasome activity, as measured by trypsin-like, chymotrypsin-like, and caspase-like activity in *skn-1* silenced worms with or without FUdR treatment. Proteasome activity is represented as slopes obtained from kinetic measurements. The experiments were conducted thrice as separate biological replicates, and significance levels (ns - not significant) were determined using an unpaired t-test with Welch’s correction.

**Figure S3.**
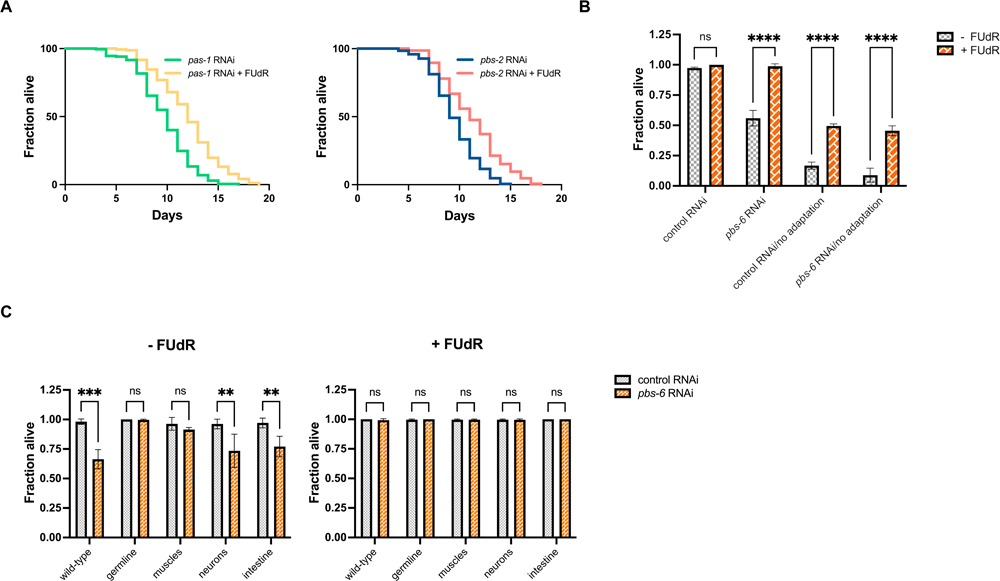
Impact of FUdR in the absence of proteasome subunits on longevity and cold survival. **(A)** The lifespan of wild-type worms when exposed to *pas-1* or *pbs-2* RNAi in the presence or absence FUdR. Number of worms used in the study: *pas-1* RNAi: n=171 (+ FUdR) or n=171 (- FUdR); *pbs-2* RNAi: n=147 (+ FUdR) or n=194 (- FUdR). **(B)** The impact of FUdR on the cold survival of wild-type and *pbs-6* knockdown worms, with or without a cold adaptation period. Data was analyzed using two-way ANOVA and the significance levels obtained from the Šidák’s multiple comparisons test are indicated for the compared conditions (ns - not significant, **** - *P* ≤ 0.0001). **(C)** The impact of FUdR on the cold survival of wild-type and *pbs-6* tissue-specific knockdown worms, with or without a cold adaptation period. Data was analyzed using two-way ANOVA and the significance levels obtained from the Šidák’s multiple comparisons test are indicated for the compared conditions (ns – not significant, ** - *P* ≤ 0.01, *** - *P* ≤ 0.001). In panels B-C, at least 90 animals were scored in three independent biological replicates.

**Figure S4.**
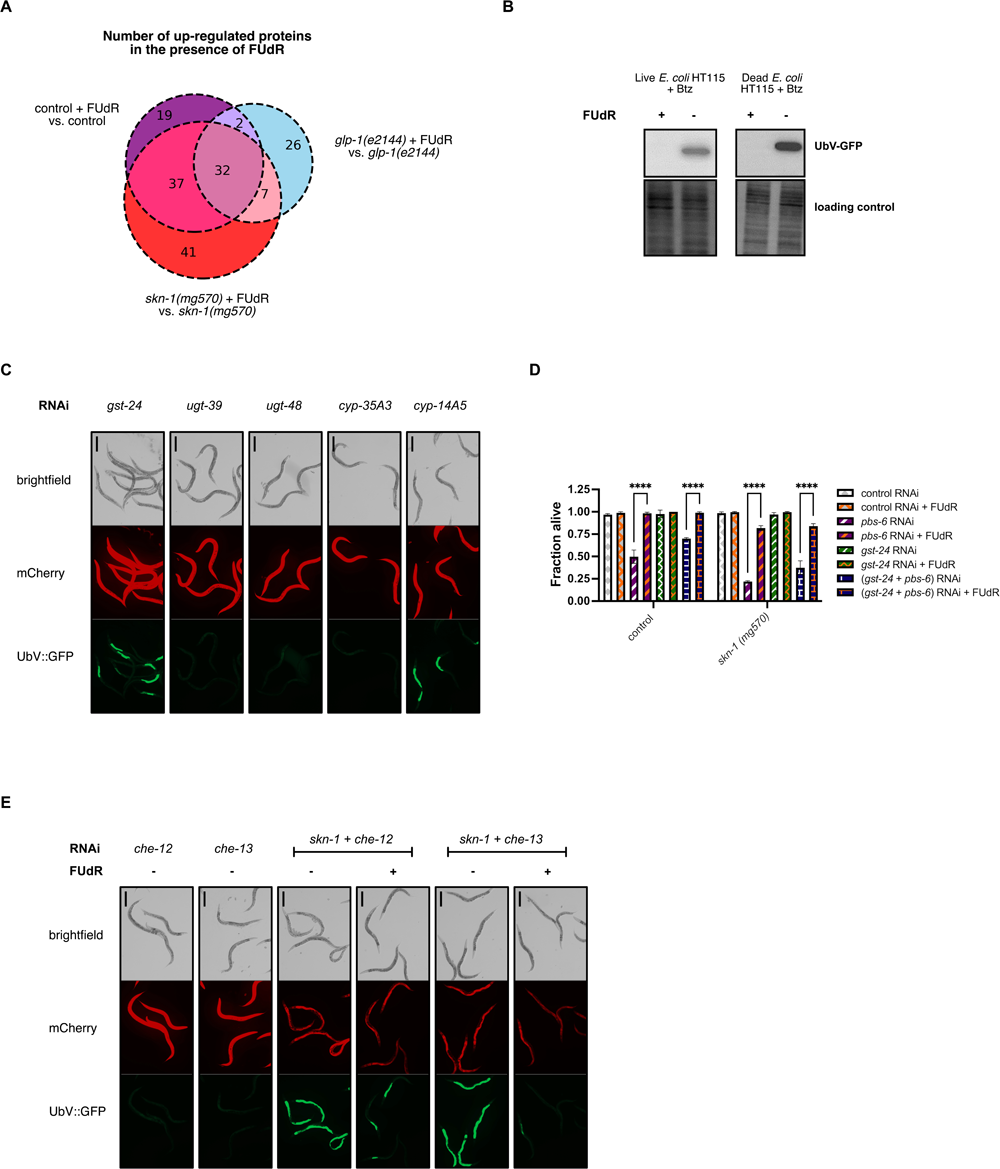
FUdR activates a detoxification pathway involving GST-24 to regulate the UPS. **(A)** Venn diagram showing the abundance of proteins up-regulated exclusively in the presence of FUdR in wild-type, *glp-1(e2144)* and *skn-1 (mg570)* worms. **(B)** Western blot showing the impact of bacterial viability on the UbV-GFP reporter turnover in the presence of bortezomib (Btz) and FUdR, as depicted by using anti-GFP antibody. The No-Stain Protein Labeling Reagent was used to confirm equal protein loading. **(C)** *In vivo* UPS activity assay showing the effect of FUdR and RNAi knockdown of detoxification-associated proteins GST-24, UGT-39, UGT-48, CYP35A3, and CYP14A5 on UbV-GFP turnover. **(D)** The impact of FUdR on the cold survival of wild-type and *skn-1(mg570)* worms subjected to control, *pbs-6* and *gst-24* RNAi. Data was analyzed using two-way ANOVA and the significance levels obtained from the Šidák’s multiple comparisons test are indicated for the compared conditions (ns - not significant, **** - *P* ≤ 0.0001). At least 90 animals were scored in three independent biological replicates. **(E)** *In vivo* UPS activity assay showing the effect of FUdR and RNAi knockdown of neuronal ciliary components *che-12* and *che-13*, either individually or in combination with *skn-1* RNAi, on UbV-GFP turnover. In panels C and E, the scale bar corresponds to 400 μm.

