## Supplementary material for "Sterility-Independent Enhancement of Proteasome Function via Floxuridine-Triggered Detoxification in *C. elegans*": Table S2

**Table S2.** Proteins up-regulated in wild-type, *glp-1(e2144)*, and *skn-1(mg570)* upon FUdR treatment with manually categorized functions.

| Protein | Function |
|---|---|
| GST-24 | Detofixication/innate immunity response |
| Y38H6C.15 | Unknown |
| Y58A7A.3 | Unknown |
| COL-41 | Constituent of cuticle |
| F08G2.5 | Unknown |
| F15D4.5 | Unknown |
| NUMR-2 | Unknown |
| FAR-3 | Lipid regulation |
| COL-176 | Constituent of cuticle |
| DUR-1 | Unknown |
| PGP-8 | Detofixication/innate immunity response |
| MAI-1 | ATP regulation |
| M01G12.9 | RNA regulation |
| PQN-22 | Unknown |
| UGT-48 | Detofixication/innate immunity response |
| Y45F10D.6 | Unknown |
| DCT-17 | Detofixication/innate immunity response |
| CYP-35A3 | Detofixication/innate immunity response |
| F54B8.4 | Detofixication/innate immunity response |
| UGT-29 | Detofixication/innate immunity response |
| DAO-2 | Unknown |
| CLEC-52 | Detofixication/innate immunity response |
| UGT-39,UGT-38 | Detofixication/innate immunity response |
| CYP-14A5 | Detofixication/innate immunity response |
| CDH-7 | Unknown |
| C14C6.5 | Detofixication/innate immunity response |
| F10D2.10 | Unknown |
| BAF-1 | DNA regulation |
| VALV-1 | Reproductive system regulation |
| SBDS-1 | Ribosome regulation |
| ZK355.8 | Unknown |
| ASD-2 | RNA regulation |

**Table S3.** Worm stages designated for specific RNAi treatments.

| **RNAi** | **Worm stage** |
|---|---|
| *pas-1* | Young adult |
| *pas-2* | Young adult |
| *pas-5* | Young adult |
| *pas-6* | Young adult |
| *pbs-2* | Young adult |
| *pbs-4* | Young adult |
| *pbs-5* | Young adult |
| *pbs-6* | Young adult |
| *rpn-1* | Young adult |
| *rpn-2* | Young adult |
| *rpn-3* | Young adult |
| *rpn-5* | Young adult |
| *rpn-6.1* | Young adult |
| *rpn-8* | Young adult |
| *rpn-9* | Young adult |
| *rpn-10* | Young adult |
| *rpn-11* | Young adult |
| *rpn-12* | Young adult |
| *rpt-1* | Young adult |
| *rpt-3* | Young adult |
| *rpt-4* | Young adult |
| *rpt-5* | Young adult |
| *rpt-6* | Young adult |
| *glp-1* | L1 |
| *skn-1* | L1 |
| *daf-16* | L1 |
| *hsf-1* | L1 |
| *pqm-1* | L1 |

| | |
|---|---|
| *gst-24* | L1 |
| *ugt-39* | L1 |
| *ugt-48* | L1 |
| *cyp-35A3* | L1 |
| *cyp-14A5* | L1 |
| *che-12* | L1 |
| *che-13* | L1 |
| *skn-1 + gst-24* | L1 |
| *skn-1 + ugt-39* | L1 |
| *skn-1 + ugt-48* | L1 |
| *skn-1 + cyp35A3* | L1 |
| *skn-1 + cyp14A5* | L1 |
| *skn-1 + che-12* | L1 |
| *skn-1 + che-13* | L1 |
| *atg-1* | Either young adults or L1 |
| *lgg-1* | Young adults |
| *fem-1* | L1 |
